# Spatial analysis of Hofbauer cell transcriptome, distribution and morphology in placentas exposed to *Plasmodium falciparum*

**DOI:** 10.1101/2023.11.27.568491

**Authors:** Ricardo Ataide, Rebecca Harding, Malindrie Dharmaratne, Yunshun Chen, Katherine Fielding, Lachlan Whitehead, Kelly L. Rogers, Casey Anttila, Ling Ling, Peter Hickey, Daniela Amann-Zalcenstein, Ernest Moya, Gomezghani Mhango, Steve Kamiza, Louise Randall, Cavan Bennett, Glory Mzembe, Martin N. Mwangi, Sabine Braat, Kamija Phiri, Sant-Rayn Pasricha

## Abstract

Placental infection remains a significant health burden for mothers and their babies in low-income countries, especially in sub-Saharan Africa, where malaria transmission is intense. An increase in inflammatory biomarkers and poor vascularisation are characteristics of placentas infected with malaria. Hofbauer cells (HBCs) – placental villous macrophages of fetal origin – are one of the most abundant immune cells in the placenta. HBCs are thought to have roles in angiogenic processes and have been linked with the pathophysiology of several infections and inflammatory conditions during pregnancy, including malaria (caused by *Plasmodium falciparum*). However, there is limited *in situ* data on the transcriptional, proteomic or morphologic profile of these cells either during or following clearance of *P. falciparum* infection. We leveraged placental samples prospectively collected at delivery from 610 Malawian women enduring a high burden of malaria and other infections and nutritional deficiencies. We profiled placentas through spatial transcriptomic and proteomic platforms to discern *in situ* HBC features that could distinguish placentas with or without evidence of past malaria. In this cohort, past placental infection was common and was associated with lower birth weight babies (adjusted effect [95% confidence interval], −80.9 [−165.9, −3.7] g, P= 0.040). However, at term, HBC numbers, abundance, and transcriptional profiles from placentas with evidence of past infection were similar to those of placentas without malaria. HBCs may recover post-infection back to a basal state or may be replaced in the tissue over the course of pregnancy. Placentas with evidence of past malaria did show evidence of reduced fetal vessel development (mean area difference: −22.8% [−37.6, −7.9], P=0.003). Reduced vascular development following infection early in pregnancy may reflect disturbances to the normal vasculogenic and angiogenic processes, of which HBCs are an integral part.

## 2. Introduction

Despite decades-long control efforts, malaria in pregnancy remains a significant public health problem, particularly in malaria-endemic African countries ^1^. Pregnant women are more susceptible to malaria infection than their non-pregnant counterparts, primarily as the placenta can become a haven for parasite strains - notably *Plasmodium falciparum* – that express novel surface variants towards placental ligands ^2^. Malaria during pregnancy, particularly placental malaria (confirmed histological evidence of parasites in the placenta)^3^, is associated with poor maternal and fetal outcomes, including severe maternal anemia, stillbirth, and low birth weight ^2,4–6^. A key strategy to reduce malaria in pregnancy, widely applied through international recommendations, is intermittent preventative treatment during pregnancy with Sulphadoxine-Pyrimethamine (IPTp-SP) beginning in the second trimester ^7^. However, this regimen misses the critical first trimester, when about 65% of first placental infections occur ^8^.

The placenta governs the crosstalk between maternal and foetal tissues and, in doing so, ensures that nutrients, metabolites, immunoglobulins, and oxygen are transferred from the mother; in addition, it removes foetal waste products and ensures that the maternal immune system tolerates the development of the foetal tissues ^9,10^. An impaired placenta is often associated with pregnancy complications ^10,11^. In the event of a *P. falciparum* infection during pregnancy, local inflammatory responses – resulting in an accumulation of maternal immune cells, particularly monocytes and neutrophils ^12–15^ in the intervillous spaces of the placenta – as well as the development of antibodies to specific parasite variant surface antigens that target placental ligands can clear the infection ^16–19^. However, this often comes at a price, with the placenta showing signs of vascular impairment, potentially as a consequence of the response mounted ^12,20–27^. Placental infection by *P. falciparum*, and to a lesser degree, *P. vivax,* leaves residual histologic evidence even after the active infection is cleared ^3,28–31^. Placental malaria can be broadly defined as active infection, if parasites are observed in the intervillous (maternal) spaces; or past infection, if the accumulation of the malaria-derived pigment haemozoin is observed within scarred chorionic villi ^3,29^, home to the resident villous macrophages, the Hofbauer cells ^32^ (HBCs).

HBCs first appear as early as day 10 of pregnancy and remain in close contact with both the maternal and the foetal blood until term ^33,34^. They are among the most numerous immune cells in the placenta ^34^ and their position in the chorionic villi, the largest area of maternal/foetal transfer, supports the hypothesis that they are key to placental – and consequently foetal – development ^35,36^. Evidence of this is the fact that these macrophages seem equipped to not only deal with infectious threats ^37^, but also to assist with vascular development ^38,39^ and to transport immunoglobins, and possibly nutrients, across the villous stroma to the foetal circulation ^40,41^. Recent advances in techniques for isolation of HBCs have allowed for single-cell RNA sequencing, which greatly increased our understanding of their potential roles ^42–45^, but lack spatial information or cell-cell interaction patterns. What does start to emerge is that HBCs are a diverse population of macrophages, or more likely, mirroring their host organ, a multipurpose population of macrophages ^46^. Several studies have investigated the role of HBCs in diabetes, viral infectious (HIV, ZIKA, SARS-CoV-2, Hepatitis), bacterial infections (Listeria) and protozoal infections (Malaria) ^35,47^; however, these studies have relied on information collected from isolated HBCs (contaminated, to different degrees, with maternal macrophages), or *ex vivo* cultured primary HBCs, lacking spatial information including the network of interactions with other placental cell types. When these cells were looked at *in situ* most studies relied on a small number of immunohistochemical markers, or a limited sample size, making it difficult to clarify their activation status and function ^48–57^. Previous studies have examined HBC response to placental infection with *P. falciparum* ^58,59^, but were limited to the scope of profiling possible. However, new opportunities are now available that enable the use of spatial transcriptomic profiling combined with multidimensional immunostaining that allow us to interrogate the tissue at unprecedented resolutions.

Since inflammation, and dysregulated angiogenesis are mechanisms proposed to drive poor outcomes during placental malaria ^12,20–27^ and HBCs are placental macrophages able to secrete inflammatory cytokines and are involved in angiogenic processes, we investigated the hypothesis that *P. falciparum* infection would modify HBCs within the placental tissue contributing to a continuing pathological process, even after clearance of maternal infection, leading to poor placental and foetal outcomes.

We leveraged placental tissue samples collected with corresponding clinical data in Southern Malawi as part of the REVAMP trial ^60^ to interrogate HBCs *in situ* through a combination of spatial transcriptomic and proteomic approaches. Our results show that, in this cohort of women receiving IPTp-SP, past malaria infection was associated with lower birth weight babies and reduced vascular development. However, at term, HBC transcriptional and morphological signatures did not differ between placentas with or without malaria infection.

## 3. Methods

### Clinical trial and placental sample collection

REVAMP was an open-label, randomised controlled trial of intravenous ferric carboxymaltose (iron) to treat antenatal anaemia that took place in the Blantyre and Zomba districts of Malawi between November 2018 and March 2021 ^60^. Malaria is highly prevalent in Malawi ^1^, with transmission occurring throughout the year. Pregnant women in their second trimester (between 13-26 weeks of gestation) were recruited if they had a capillary haemoglobin of < 10 g/dL (i.e., moderate or severe anaemia) and a negative malaria rapid diagnostic test (mRDT, SD Bioline Malaria AG Pf/Pan, Standard Diagnostics). Women presenting with a positive mRDT at pre-screening were given anti-malaria treatment with arthemeter-lumefantrine and allowed to enter the trial if they were malaria-negative by microscopy a week after initiating the treatment and met the remaining inclusion criteria ^60^. At enrolment, 862 participants were randomised to receive a single infusion of ferric carboxymaltose at study entry or a full course of twice daily oral iron (standard of care) for the remainder of pregnancy. Also at enrolment, all women received directly observed intermittent preventive malaria treatment with Sulphadoxine-Pyrimethamine (IPTp-SP) – unless contraindicated (e.g. recent malaria treatment, or HIV positive taking Cotrimoxazole). Demographic and clinical variables together with biological samples (including venous blood) were collected throughout pregnancy and at delivery ^60,61^.

At delivery, a trial team nurse assisted with the delivery and the clinical management of the participant and their babies. The placenta was collected and kept in PBS until processing in the labour ward. Two 2cm x 2cm x 1cm biopsies were taken from an off-centre position, halfway between the umbilical cord and the edge of the placenta. Care was taken to avoid lesions, areas of tissue tear, missing material or infarcted areas. The biopsies were placed in 50 mL of 10% buffered formalin, taken to the laboratory in Zomba, Malawi and stored for 24h at 4°C, after which the formalin was changed. Within one week the tissues were transferred to 80% ethanol until further processing. The samples were paraffin-embedded, using standard techniques, and one 5 μm-thick section stained with Haematoxylin-Eosin (H&E) remained in Malawi for histopathology evaluation. All the formalin-fixed paraffin-embedded (FFPE) blocks were shipped to the Walter and Eliza Hall Institute, Melbourne, Australia.

### Histopathology evaluation of placental malaria status

H&E slides of full-thickness placental sections were evaluated independently in Malawi and in Australia by trained histopathologists, each with >5 years’ experience in placental malaria evaluation. Where discrepancies were found, the microscopist in Australia reviewed the discrepant slides, blinded to the initial reports, and that third reading was used to resolve the discrepancies. To evaluate placental malaria status the microscopists reported on the presence/absence of malaria parasites and/or malaria-derived haemozoin in the intervillous spaces of the placenta, as well as in the placental tissues. Haemozoin was confirmed and distinguished from formalin-pigment by reading the slides under polarised light. Placentas were classified as Absent if no evidence of current or past *Plasmodium* infection was detected; Active if *Plasmodium* parasites were detected in the intervillous spaces of the placenta and; Past if haemozoin was detected in the tissues, in the absence of parasites in the intervillous space (adapted from ^3^).

### Tissue microarrays

Regions of chorionic villi were manually identified on H&E slides. Five different tissue microaarays (TMAs) were built. Two TMAs consisted of 18 0.5 cm x 0.5 cm tissue sections to be used for spatial transcriptomics, and three TMAs containing a total of 126 2 mm diameter cores to be used for Opal multiplexed immunofluorescence. All TMAs were sectioned and stained with H&E and Perls (for tissue quality, architecture and malaria evaluation/confirmation). For the spatial transcriptomics TMAs, placentas were chosen from the oral-iron arm of the trial. For the Visium 10X experiments, one placenta belonging to each malaria group was randomly chosen out of the 18 tissues. For GeoMX, all 18 tissues were evaluated. For the immunofluorescence TMAs, the number of placentas chosen was based on preliminary experiments on 12 tissues that determined 42 placentas per group would be required for an 80% power to differentiate between Absent and Past placentas based on proportion of cells expressing HIF1 (alpha = 0.05, m1 = 0.5, m2 = 1.12, sd = 1) and 3 placentas per group to differentiate Absent and Active placentas (alpha = 0.05, m1 = 0.5, m2 = 4.0, sd = 1).

### Spatial transcriptomics – Visium slide preparation and sequencing

Briefly, Visium Spatial for FFPE Gene Expression Kit, Human Transcriptome, 4 reactions (#1000337, 10x Genomics) were used according to the manufacturer’s recommendations. Each area contains ~5,000 detection spots of 55 µm in diameter and a 100 µm centre-to-centre distance. Each slide included a placental sample purchased from Cureline, Inc., CA, USA to serve as a control for the quality of the tissue fixation and processing conducted in Malawi and to provide an inter-reliability control for the Visium reactions (Supplementary Figure 1). Gene expression libraries were pooled and sequenced on the Illumina NextSeq2000 according to 10X Genomics’ recommendations. Illumina output from 10x Visium sequencing was processed using spaceranger 2.0.0. Downstream analysis of the Visium data were restricted to spots within the placental villi that were enriched for HBCs, defined as spots with non-zero *LYVE1* counts. We used the expression of *LYVE1*, as it is a unique marker of HBCs within cell populations from the chorionic villi (representative data from CellxGene data from Vento-Tormo et al. - https://maternal-fetal-interface.cellgeni.sanger.ac.uk/ -, Figure 2B. Lowly expressed genes were filtered using the filterByExpr function in edgeR (v3.41.3) followed by TMM normalisation and a quasi-likelihood pipeline was used to identify differentially expressed genes between absent, past and active malaria groups.

**Figure 1 –.**
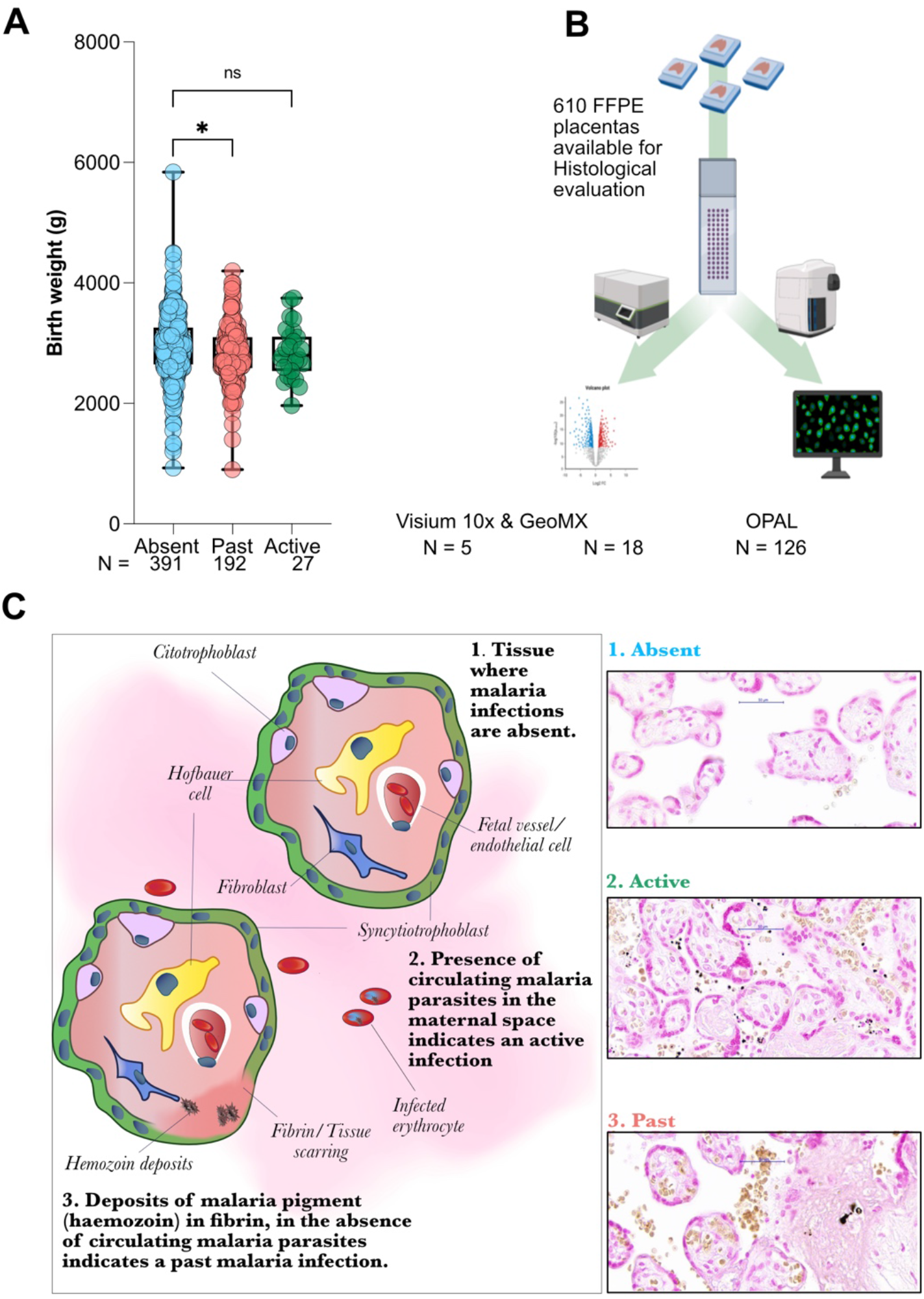
Clinical outcomes of past malaria, techniques and definitions. (A) Birth weight of babies from whom a placental sample had adequate tissue quality and a birthweight recorded from the REVAMP trial according to placental malaria status (N = 610). * adjusted P = 0.040, ns – not significant. (B) Six hundred and ten placentas in FFPE blocks were evaluated by histopathology for placental malaria. Tissue microarrays were constructed for Genomics 10x Visium (N=5), Nanostring GeoMX (N=18) and OPAL Polaris multiplexed immunostaining (N=126). (C) Illustration and representative Perls-stained slides representing placental malaria status, as defined in this study. Placentas were evaluated for malaria status using H&E-stained slides under polarised light. TMAs were additionally stained with Perls stain to enhance the presence of iron-containing haemozoin within scarred tissue.

**Figure 2 –.**
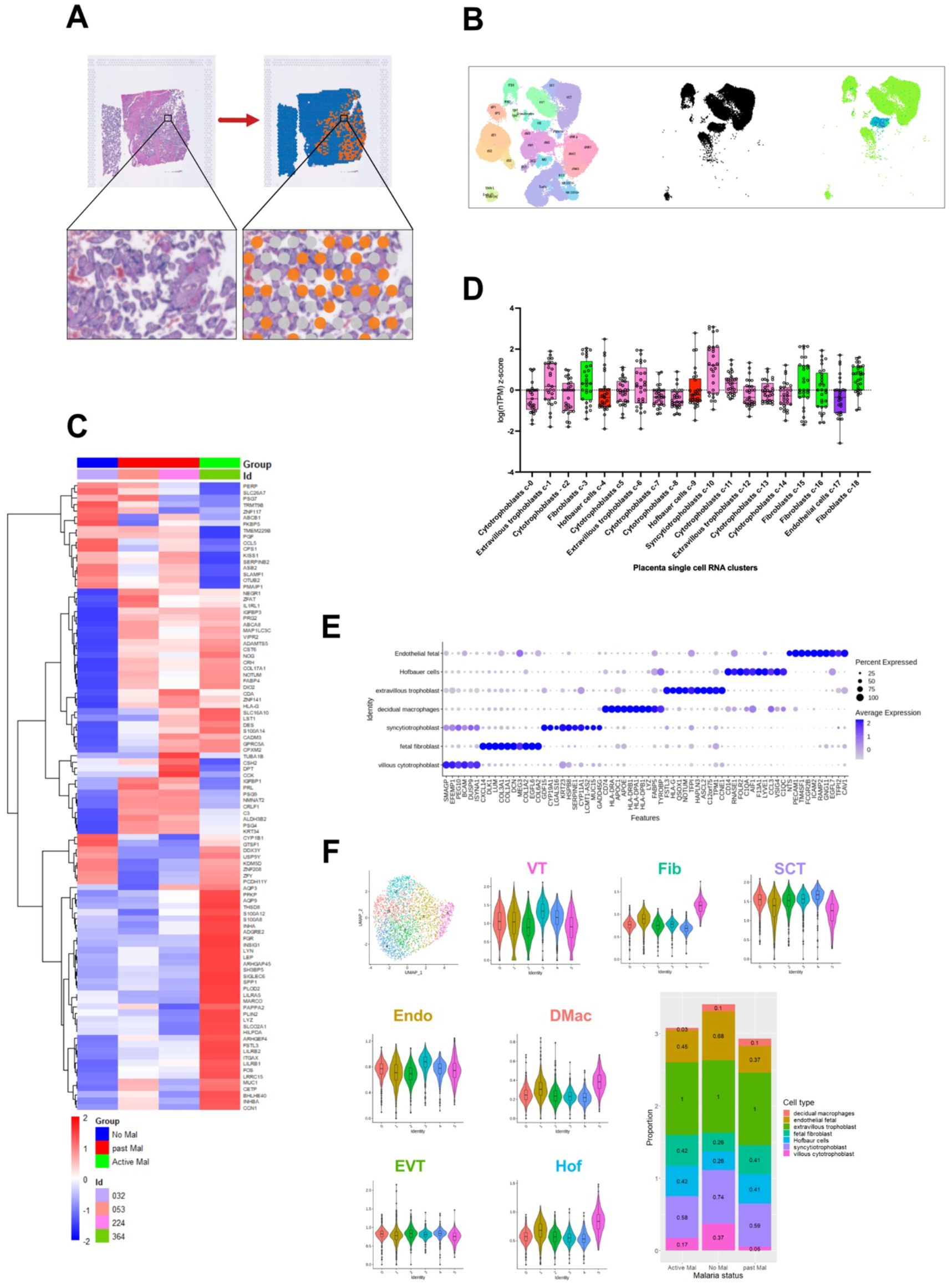
Genomics 10x evaluation of placentas with or without evidence of malaria infection. (A) One reaction chamber of a Visium reaction slide with placental tissue visualised in Loupe Browser 6. Areas of chorionic villi were manually curated before selecting HBC containing spots for analysis (*LYVE1*-positive spots, highlighted in orange). (B) Single-cell RNA sequencing data from Vento-Tormo et.al.^42^ publicly available through the CellxGene website (https://maternal-fetal-interface.cellgeni.sanger.ac.uk/) demonstrating the specificity of *LYVE1* to identify HBCs within the placenta. The left hand panel reflects all cells isolated belonging to the Maternal blood, the foetus or the placenta. The centre panel, in black, identifies only those cell subsets identified as Placenta. The right-hand panel shows *LYVE1* expression (blue) exclusively within the HBC cluster. (C) Heatmap of the overall differential transcription of the 100 highest expressed genes across all tissues between placentas were malaria was absent (n=1), active (n=1) or past (n=2). (D) Evaluation of the cell types most often represented within the Visium spots with detected *LYVE1* transcription. We took the 30 most expressed genes found in this study and we extracted the z-score of the log (number of transcripts per million) obtained when these genes were mapped to the clusters identified by Vento-Tormo et.al. ^42^. Trophoblast clusters (represented in pink) and Fibroblast clusters (green) were predominant. (E) Deconvolution strategy employed to try and achieve a higher resolution for the cell-types represented in *LYVE1* Visium spots. The top 10 marker genes for each cell type in the reference data by Vento-Tormo et.al. (F) The spatial transcriptomic data of all samples were integrated and clustered. Gene signature scores were calculated on the integrated data using gene signatures derived from the reference data. HBCs and Fibroblasts were highly represented in cluster 5. The bar plot depicts the proportion of spots expressing the seven main cell types from the reference data in active malaria, no malaria and past malaria.

### Spatial transcriptomics – GeoMx slide preparation and sequencing

TMAs of 18 placental tissues (6 each with past, active or absent placental malaria) were serially sectioned. Consecutive sections were either stained for H&E or sent to Nanostring, Seattle, where the sequencing was performed. Four morphology markers (PanCK - Syncytiotrophoblast, CD163 - HBC, CD31 – fetal vessels, and DAPI - nuclei) were used to identify tissue architecture and guide the investigator-led selection of regions of interest (ROIs) (see example in Figure 3A). Within each ROI, a mask was applied to CD163+ targets generating segments from which whole transcriptome information was collected (Figure 3A). All segments generated reached sequencing saturation. A total of 18,676 targets were identified in the samples, with 8,703 genes expressed in 10% of the segments and 2,470 genes expressed in 50% of the segments (Supplementary Figure 2). Genes with a total count of less than 10 were filtered from the analysis, followed by TMM normalisation.

**Figure 3 –.**
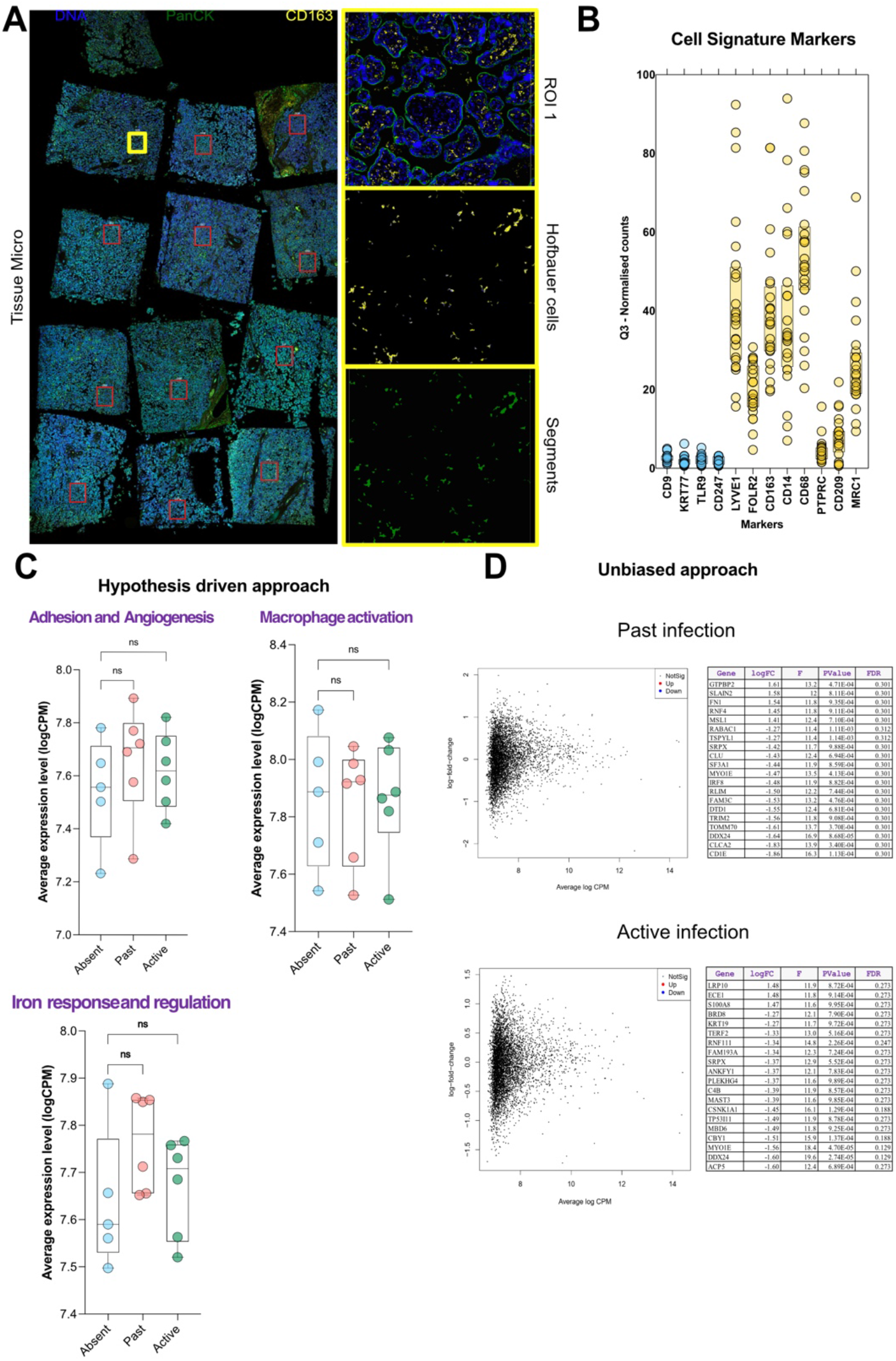
NanoString GeoMX™ evaluation of placentas with or without evidence of placental malaria. (A) FFPE TMAs were stained for the morphology markers DAPI (cell nuclei), PanCK (SCT), CD31 (fetal vessels), and CD163 (HBCs). After staining, one 700 μm wide ROI was applied to each of the 18 tissues. The Rare Cell Profile approach was used manually select villi – to exclude intervillous areas – and after applying a CD163 mask, segments were generated from which only those DNA oligos bound to HBCs were collected for sequencing. (B) Hallmark HBC genes (in yellow) were highly represented after sequencing. Results are given as counts after Q3-normalization. Genes in blue represent examples of genes associated with the SCT, cytotrophoblasts, fibroblasts and endothelial cells. (C) Hypothesis-driven approach to differential gene activation. Genes belonging to 3 distinct pathways (gene tables in Supplementary figure 5) were analysed for their differential transcription levels between placentas without evidence of malaria (Absent - blue), with past malaria (Past - red) or with an active infection (Active - green). (D) Unbiased approach to differential gene activation. MD plots of the differential transcription of genes between malaria Past and Absent placentas (top) and Active and Absent placentas (bottom) are shown. No genes were found to be either up-regulated or down-regulated after adjustment for false discovery. The top 20 genes, based on logFold change are listed in the adjacent tables. * One-Way Anova adjusted P < 0.05; ** One-Way Anova adjusted P < 0.01; ns – not significant.

### Visium data deconvolution

Deconvolution of the Visium spatial transcriptomics data was performed using scRNA-seq data from Vento-Tormo et al. ^42^ as the reference. First, the Visium samples were integrated using the Seurat (v4.2.1) integration pipeline for scRNA-seq data where spots were treated as cells. The integrated data was clustered using the FindClusters function in Seurat with resolution parameter 0.9. Cell type specific signature genes for the reference data were identified using the FindAllMarkers function from Seurat with the following threshold parameters: min.pct = 0.5, logfc.threshold = 2, and min.diff.pct = 0.25. These signature genes were then used to calculate gene signature scores on the integrated Visium data. The gene signature scores were calculated as the average gene expression of the set of signature genes for each cell type. The gene signature scores were overlaid on the clusters to investigate if any of the six identified clusters were enriched for known cell types from the reference data.

### Chromogenic immunohistochemistry and Opal Polaris multiplexed immunofluorescence

All antibodies used for multiplexed immunofluorescence were validated individually first by chromogenic singleplex immunohistochemistry (IHC) and then by singleplex immunofluorescence using the Opal 7 kit reagents (NEL871001KT, Akoya Biosciences, Waltham, MA, USA). Detailed methods and antibody information are in supplementary data. For Opal multiplex immunostaining, 2mm core TMAs were prepared after manual selection of areas of Chorionic villi. Opal staining followed standard Opal-recommended protocols. All slides were scanned on a PhenoImager-HT with a 0.45 NA objective.

### Immunofluorescence analysis of spatial and morphological features

Multiplex image analysis was performed using a custom pipeline in FIJI ^62^. For each placental tissue sample – consisting of a 2 mm core on a TMA – 3 regions of chorionic villi were manually selected, cropped, and analysed independently. Placental tissue and intervillous space, as well as foetal vessels and tissue-associated cells were segmented with a trained ilastik ^63^ pixel classifier. Nuclei were segmented with a custom trained Stardist network ^64,65^. Cells were then scored based on manually defined thresholds for Opal stains, and cell edge - tissue edge distances measured for each positive cell. Median intensities were also recorded and mask images produced to allow quality for control checking by the researchers. Results for each region were pulled and averaged for each placental tissue before the final analysis. Instances where the quality of the staining, the tissue or the annotation did not pass the quality control checking resulted in those tissues being dropped from the analysis. The numbers of tissues dropped are reported next to each of the analysis.

### Statistical analysis

All placental tissues with adequate tissue quality, and where two independent histological evaluations were performed, were included in the analysis of clinical data. Analysis of transcriptomic and proteomic data consisted of a subset of samples as described above. The association between placental malaria and birth weight was estimated using unadjusted and adjusted linear regression models, with the adjustment set determined a priori and consisting of gravidity, gestational age at delivery, infant sex and trial arm to obtain the direct effect. Similar associations between placental malaria and characteristics of the placenta were explored. No adjustment for multiplicity was used, instead estimates and confidence intervals alongside p-values were reported to support the reader’s interpretation of the findings. Group comparisons were achieved after regression across groups, either unadjusted or adjusted for gravidity, maternal HIV status, maternal anaemia at delivery, maternal iron-deficiency at delivery, gestational age at delivery, infant sex and trial arm. All statistical modelling was performed using Stata/SE 17.0 (StataCorp, TX, USA). Graphical visualisation was done using GraphPad Prism 9.0 (GraphPad Software, LLC) and panels were prepared in Affinity Publisher 2.1.1 (Pantone, LLC).

### Ethics

Ethics for this work were obtained from the College of Medicine Research Ethics Committee (P.02/18/2357) and the WEHI Human Ethics Research Committee (18/02). The trial was approved by the Malawian Pharmacy and Medicines Regulatory authority (PMRA/CTCR/III/25052018100). Informed consent was obtained from all participants for collection, transport and use of the samples.

## 4. Results

### Past malaria is associated with lower birth weight

Almost 7% of the >20,000 women screened for inclusion in the REVAMP trial were mRDT positive^60^. The REVAMP trial enrolled 862 second-trimester pregnant women, from whom 710 placentas were collected. Of those, 610 placentas were from singleton pregnancies and had adequate tissue quality – fixation and paraffin-embedding – that allowed for two independent evaluations of histopathology of current or past malaria. Twenty-nine (4.8%) placental samples showed active infection (20 with both active and past infection – commonly designated as chronic infections – and 9 with active infection only). One hundred and ninety-three (31.6%) placental samples showed past malaria with no evidence of circulating parasites. Placental malaria (past and active) was associated with lower birth weight (mean difference [95% confidence interval], p-value: −119.3 g [−202.5, −36.1], P = 0.005). This was evident in cases exhibiting past placental malaria compared to those without malaria infection, after adjusting the model for gravidity, maternal HIV status, maternal anaemia at delivery, maternal iron-deficiency at delivery, gestational age at delivery, infant sex and trial arm (−80-9 g [−165.9, −3.7], P = 0.040) (Supplementary table 1).

**Table 1 –.** Clinical and demographic characteristics of the participants with placental tissues available.

|  | All participants* | Absent (no malaria)<sup>a</sup> | Past malaria<sup>b</sup> | Active malaria<sup>c</sup> |
| --- | --- | --- | --- | --- |
| N (%) | 610 (100) | 388 (63.9) | 193 (31.6) | 29 (4.8) |
| Trial arm (%) |  |  |  |  |
| Oral iron | 49.6 | 47.7 | 51.8 | 51.7 |
| IV iron | 50.4 | 52.3 | 48.2 | 48.3 |
| Maternal age at enrolment, years, median [IQR] | 20.0 [18.0-25.0] | 22.0 [19.0-28.0] | 18.0 [17.0-20.0] | 19.0 [17.0-21.0] |
| Maternal gravidity, median [IQR] | 1 [1-2] | 2 [1-3] | 1 [1-2] | 1 [1-2] |
| Primigravidae (%) | 53.6 | 42.0 | 74.1 | 74.4 |
| Secundigravidae (%) | 22.8 | 26.8 | 16.1 | 13.8 |
| Multigravidae (%) | 23.6 | 31.2 | 9.8 | 13.8 |
| Maternal weight at enrolment, Kg, mean (SD) | 55.4 (8.2) | 55.9 (8.6) | 54.4 (7.6) | 54.1 (6.5) |
| Maternal GA at enrolment, weeks, mean (SD) | 21.5 (3.3) | 21.3 (23.4) | 21.9 (2.8) | 21.3 (3.3) |
| Maternal GA at delivery, weeks, mean (SD)<sup>§</sup> | 39.5 (2.1) | 39.5 (2.1) | 39.4 (2.1) | 39.1 (1.6) |
| Maternal HIV status (%)<sup>§§</sup> |  |  |  |  |
| Positive | 16.4 | 21.6 | 6.8 | 10.3 |
| Negative | 83.6 | 78.4 | 93.2 | 89.7 |
| Maternal anaemia at delivery (%)<sup>§§§</sup> |  |  |  |  |
| Yes | 29.4 | 31.1 | 25.6 | 37.9 |
| No | 70.6 | 68.9 | 74.4 | 62.1 |
| Maternal iron-deficiency at delivery (%)<sup>§§§§</sup> |  |  |  |  |
| Yes | 30.5 | 38.7 | 17.0 | 10.3 |
| No | 69.5 | 61.3 | 83.0 | 89.7 |
| Infant parameters: |  |  |  |  |
| Live birth (%) | 99.0 | 98.7 | 99.5 | 100.0 |
| Birth weight, g, mean (SD)<sup>d, †</sup> | 2894.7 | 2937.8 | 2814.8 | 2844.8 |
|  | (503.8) | (521.5) | (467.3) | (432.5) |
| Birth length, cm, mean (SD) <sup>††</sup> | 47.6 (3.4) | 47.9 (3.5) | 47.3 (3.5) | 46.6 (3.1) |
| Infant sex (%) <sup>†††</sup> |  |  |  |  |
| Male | 50.2 | 47.8 | 53.7 | 59.3 |
| Female | 48.8 | 52.2 | 46.4 | 40.7 |

N – Number; IV – Intravenous; IQR – interquartile range; SD – standard deviation GA- Gestational Age. \* Placental samples are from women who delivered a singleton liveborn or stillborn infant.
<sup>a</sup> No malaria is defined as absence of tissue evidence for malaria (parasites or pigment).
<sup>b</sup> Past malaria includes only those women where malaria pigment was detected in the placenta tissue in the absence of any circulating parasites.
<sup>c</sup> Active malaria includes tissues where we detected the presence of circulating parasites in the intervillous space. Twenty placental tissues had evidence of pigment in the tissue in addition to circulating parasites, and thus could have been further classified as Chronic infections<sup>3</sup>.
<sup>d</sup> Results for unadjusted and adjusted analysis are presented in supplementary table 1.
<sup>§</sup> Eleven participants were missing a recorded gestational age at delivery (8 in the Absent group, and 3 in the Past group).
<sup>§§</sup> Five participants were missing a recorded HIV status (4 in the Absent group, and 1 in the Past group).
<sup>§§§</sup> Forty-one participants were missing an Anaemia status at delivery (28 in the Absent group and 13 in the Past group). Anaemia is defined as a venous haemoglobin of <110 g/L.
<sup>§§§§</sup> Sixteen participants were missing an iron-deficiency status at delivery (11 in the Absent group and 5 in the Past group). Iron-deficiency was defined as a venous ferritin < 15 mg/L, adjusted for CRP.

### Spatial transcriptional profiles of placental tissue with and without malaria infection

We hypothesised that HBCs would exhibit a state of inflammation in the tissue that would persist beyond resolution of active infection. Overall comparison of gene expression in *LYVE1*-enriched spots across tissues revealed several gene sets differentially expressed between malaria infected (past or active) and malaria-absent tissue, as seen after analysing the differential expression of the 100 most highly expressed genes across tissues (Figure 1C). Pathway analysis revealed up-regulation in Cell Adhesion, Neuroactive ligand-receptor interactions, and Kaposi sarcoma-associated herpesvirus pathways, as well as a down-regulation of ABC Transporters associated with malaria infection (Supplementary figure 3). Interestingly, the two tissues with evidence of past malaria had highly similar transcriptional profiles (Figure 1C). To try and identify the main cell types contributing to those responses, we mapped the transcript abundance of the top-30 expressed genes onto the 18 clusters identified by Vento-Tormo et al ^42^. The number of Transcripts per Million (represented as a z-score to facilitate comparisons between genes) of each of the 30 genes was plotted, revealing that our *LYVE1*-enriched spots had genes that were more often associated with syncytiotrophoblast (SCT) and cytotrophoblast (CTB) as well as fibroblasts (Figure 2D). This is aligned with the tissue architecture of placental villi. By using stringent cut-offs for identifying differentially expressed genes in the reference data, we curated the gene sets that would better differentiate the most common cell types within the placenta (Figure 2E). These gene signatures were used for cell type deconvolution of the Visium spots. First, we performed integration of the spatial transcriptomic data from all samples following Seurat’s integration pipeline for scRNA-seq data. Following cluster analysis of the integrated data, and we identified six main clusters of spots. Overlaying the gene signatures from the reference data on to the integrated data revealed cluster five to be enriched in HBC and fibroblast gene signatures.

### Hofbauer cell-specific transcription profiles

After evaluating the tissue-wide transcription profile of placentas – with a focus on areas of expression profiles consistent with HBCs – we next evaluated the specific transcriptional profile of HBCs *in situ* with Nanostring’s GeoMX platform, using the Rare Cell profiling approach. Markers of HBCs were highly enriched in our sample compared to genes that identify most other chorionic villi cell types (Figure 3B, Supplementary figure 4). An analysis using the same combination of gene signatures used for the Visium deconvolution approach revealed very similar results, with fibroblast signatures also being strongly represented (Supplementary figure 4). We hypothesised that genes involved in Adhesion and Angiogenesis, Macrophage Activation and Iron Response and Regulation pathways would be differentially expressed between placentas with or without evidence of past malaria but did not find evidence for differential expression (Figure 3C and supplementary figure 5). Similarly, after adjustment for false discovery, there was no significant difference between overall differential expression in HBCs between placentas with active or past malaria and those without evidence of malaria infection (Figure 3D).

### Morphological and spatial features of Hofbauer cells

We next investigated the spatial distribution and morphology of HBCs. TMAs comprising cores of chorionic villi with a 2mm diameter (N=126 cores) were constructed (example in Figure 4A). We quantified parameters related to tissue architecture (Figure 4B), as well as parameters related to HBC morphology and abundance (Figure 4C), activation (Figure 4D), and tissue location (Figure 4E).

**Figure 4 –.**
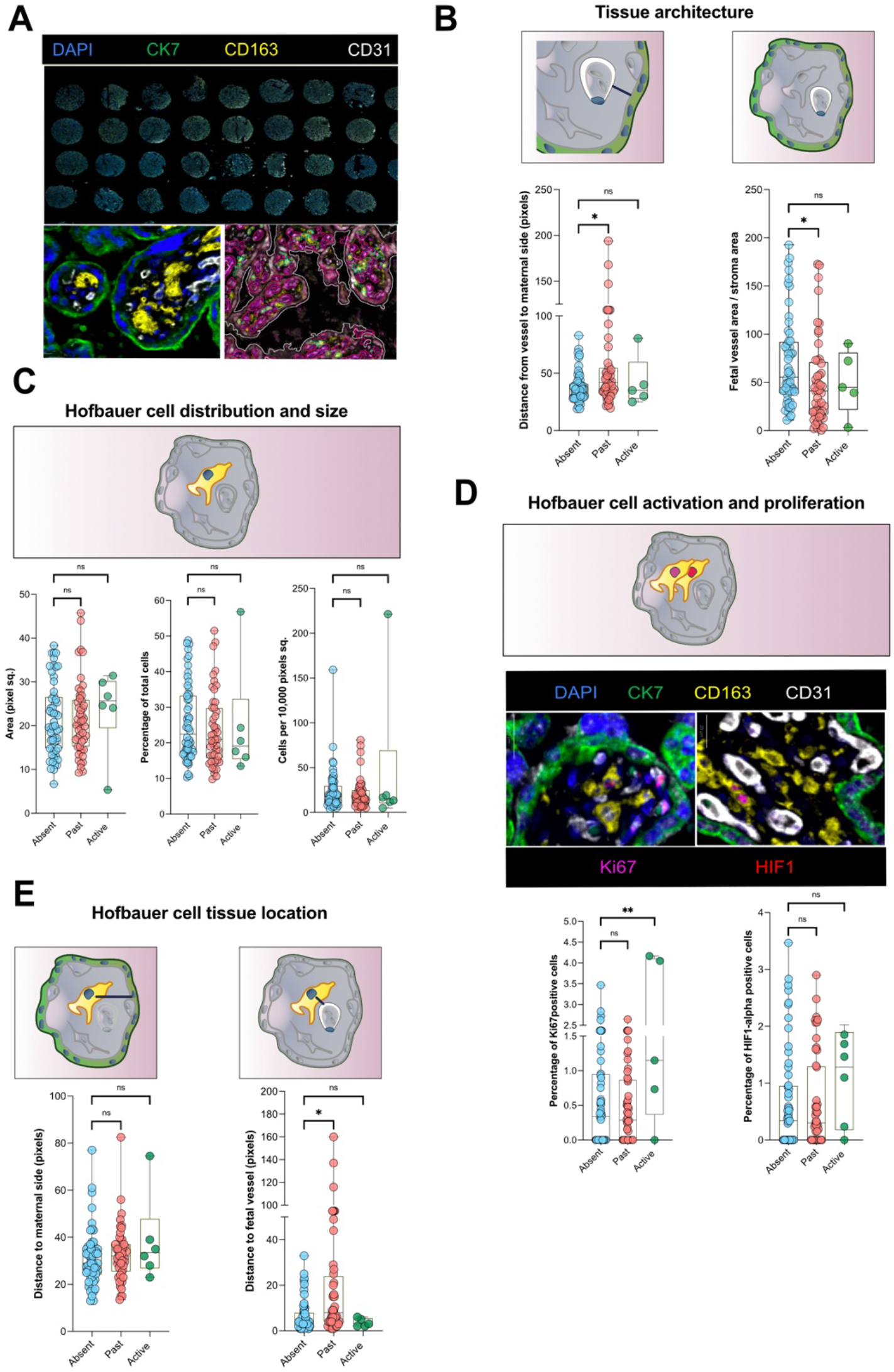
Akoya Opal Polaris multiplexed immunostaining of placentas with or without evidence of malaria infection. (A) Areas of chorionic villi from placentas in FFPE were selected to build 3xTMAs with 2-mm cores. TMAs were sectioned and stained using with Opal Polaris 7-colour kits: DAPI (blue), CK7 (green), CD31 (white), CD163 (yellow), HIF1-alpha (red), Ki67 (magenta) and HLA-DRA (orange, data not shown). A high magnification of a section of chorionic villi (40x) demonstrates the staining. An example of the result of this algorithm is also shown. (B) Fetal vessel parameters. (C) HBC distribution and morphology (D) HBC proliferation and activation. (D) HBC spatial parameters; distance to SCT and distance to foetal vessels. * adjusted P < 0.05; ** adjusted P < 0.01; ns – not significant.

Evaluation of the tissue architecture revealed a smaller percentage of the villi occupied by fetal vessels – identified as CD31-positive areas within the tissue – in placentas with past malaria when compared with those with absence of malaria (mean difference [95% confidence interval], −22.8% [−37.6, −7.9], P=0.003) (Figure 4B). This association was maintained after adjusting for gravidity, maternal HIV status, maternal anemia at delivery, maternal iron-deficiency at delivery, gestational age at delivery, infant sex and trial arm (−18.3% [−36.6, −0.1], adjusted P=0.049) (Supplementary table 2).

HBCs were not significantly different in terms of their abundance in the tissue relative to all other cells (one-way ANOVA adjusted P = 0.657), their tissue density (adjusted P = 0.07) or their individual size (adjusted P = 0.829) when compared between placentas with or without evidence of past or active malaria (Figure 4C). We evaluated HBC proliferation status using Ki67 and found a significantly higher percentage of HBC expressing Ki67 in the active group compared to placentas where malaria was absent (mean difference: 1.4 [0.5, 2.3] %, adjusted P = 0.002) (Figure 4D, Supplementary table 2) – although, overall, the percentage of cells positive for Ki67 was low (mean % Ki67 (SD) in Active = 1.7 (1.9)%). This observation was modelled with adjustment for gravidity, maternal HIV status, maternal anemia at delivery, maternal iron-deficiency at delivery, gestational age at delivery, infant sex and trial arm. The activation marker HIF1-alpha – also a hypoxia marker – did not reveal differential expression across malaria groups (adjusted P=0.154).

Relative to placentas without malaria, HBCs from placentas with evidence of past malaria were further from foetal vessels (mean difference: 15.8 [3.3, 28.4] pixels, adjusted P = 0.014) (Figure 4D, Supplementary table 2) but showed a stable position in the tissue relative to the SCT (1.2 pixels, [−4.4, 6.8], adjusted P = 0.667). This observation was modelled with adjustment for gravidity, maternal HIV status, maternal anemia at delivery, maternal iron-deficiency at delivery, gestational age at delivery, infant sex and trial arm.

## 5. Discussion

We successfully applied *in situ* transcriptome-wide and cell-specific spatial gene expression profile techniques, together with multiplexed immunohistochemistry approaches, to a cohort of placental samples collected at delivery in a setting where malaria infection is intense. We found that, in Malawian pregnant women receiving malaria prophylaxis from the second trimester onward, and where past malaria infection is associated with lower birth weight babies and reduced fetal vasculature, HBCs in placentas with evidence of past malaria do not significantly differ transcriptionally or proteomically from those where there is no evidence of placental infection.

Although both inflammation and parasitemia – resulting from maternal infections with *P. falciparum* during pregnancy – can be cleared, placentas infected with *P. falciparum* have been shown to have pigment deposition, tissue scarring and reduced vascular development (likely as the result of inflammatory processes), even in the presence of IPTp-SP^3,20–22,29^. We hypothesised that the clearance of maternal infection and inflammation would leave unaddressed inflammatory processes within the placenta tissue that would contribute to a perpetuating pathological state. HBCs – chorionic macrophages capable of responding to external stimuli and with a role in angiogenic processes in the placenta – provided a likely candidate that could respond to this tissue insult and perpetuate the pathological process throughout pregnancy, contributing to low birth weight. Our results indicate this is not the case.

To the best of our knowledge, our study is the first to have successfully applied spatial transcriptomic technology to profile the transcriptional status of HBCs *in situ* in placentas from settings where malaria and other infections are endemic. We used two different transcriptomic approaches to determine the differential expression of genes between HBCs in placentas with or without evidence of malaria infection, and our analyses highlights the opportunities and limitations of these emerging technologies. The 10X Genomics Visium platform identified clear differences in overall gene expression between placentas with or without evidence of malaria infection. A limitation of the platform is the large capture areas – spots are 55μm diameter with 100μm spacing – meaning it could not offer the near single cell resolution we needed for HBC analysis and hence we did not pursue it with a higher number of samples. We developed strategies that can overcome this limitation by leveraging published scRNA-seq data from human placentas ^42^ to generate unique cell-signature markers and deconvolute the data obtained to identify clusters of transcription where HBCs were enriched. Through this approach we identified malaria-infected placentas as having higher HBC and fibroblast transcriptional signatures, as well as lower levels of SCT-related transcription. Despite this limitation, Visium 10X may ultimately offer value as a technique to evaluate the overall transcription profiles of FFPE placental tissue exposed to varied conditions.

We took a more targeted approach to directly profile HBC gene expression profiling with the Nanostring GeoMX platform, making use of their Rare Cell profiling tools. We did not observe any significant difference in the transcription of either manually curated gene sets or unbiased overall gene expression between HBCs in placentas with or without evidence of malaria. This finding, using a technique that more reliably targets HBC *in situ* transcription, provides evidence that CD163+ HBCs in placentas with past malaria, at term, do not represent a distinct population of cells to CD163+ HBCs present in placentas without malaria.

Isolated primary HBCs can be stimulated to respond to external antigens ^37,47,58,66^, but it is unknown whether these cells are then able to return to a basal state post-stimulation. Another interesting possibility is the ability that HBCs may have to transition to myofibroblasts. The existence of macrophage-to-myofibroblast transition has recently been demonstrated in tissues such as the kidney ^67,68^, the retina ^69^, and even the chorioamnion in the placenta ^70^. Analysis of the data from GeoMX, using the Visium deconvolution gene sets, did not reveal an increased fibroblast signature in placentas with past malaria relative to those without (Supplementary figure 3). However, the impact of such a transition occurring in the chorionic villi of placentas where inflammation occurs, and the contribution it may make to the ‘disappearance’ of inflammatory Hofbauer cells from the tissues, still needs to be evaluated. What we can say is that, together with the paucity of trophoblastic and foetal endothelial signatures in the GeoMX data, the consistent appearance of a fibroblast signal indicates that HBCs form closer associations with fibroblasts than with any other placental cell type at term.

*In* situ proteomic analysis of HBCs in the context of disease has usually been performed on a limited number of tissues, using only one or two colours (thus failing to properly contextualise these cells in the tissue) and often relying on subjective methods of quantification. We overcame these limitations by conducting multiplexed immunostaining on manually selected chorionic villus areas from 126 placental samples, and performing automated algorithm-driven image analysis. Studies have recently evaluated the role that pathogens play in activating HBCs and promoting a proliferative response (reviewed in ^34,47^). We thus looked at the expression levels of the markers Ki67 (proliferation) and HIF1-alpha (activation). Although tissue macrophages are recognised to have self-proliferative capacity ^71,72^, this feature in HBCs remains uncertain ^71^. A scRNAseq study suggested that HBCs have self-proliferating capability ^42^, and studies of ZIKA-infected placentas were able to show some increased HBC proliferation in the tissues marked by the expression of Ki67 ^51,73^, suggesting an ability to respond to external pathogens. In our cohort of samples, the overall proportion of cells expressing these markers, was very low. The transient, but not sustained, increase in Ki67 expression in HBCs from infected placentas indicate that HBCs may indeed respond to infection with increased proliferation and expression of activation markers but then either return to a basal state, or be replaced in the tissue.

As pregnancies progress to term, foetal vessels within chorionic villi expand and occupy a larger proportion of the villus area, to a point where, in terminal villi, little separates the foetal and maternal circulations ^11,74^. Placental malaria infections may disrupt the vascular development, and this is most commonly observed as abnormal levels of proteins on the angiogenic pathway ^22,24,25,75,76^ or as actual physical changes to the architecture of the placental tissue ^31^. In line with these observations, placentas in our cohort with evidence of past malaria had a lower proportion of foetal vessel area (measured as CD31+ areas) than placentas where no infection was detected, suggesting a potential effect of *P. falciparum* on the development of the foetal vasculature early in pregnancy. This larger area of stroma in placentas with evidence of past malaria resulted in HBCs being positioned further away from foetal vessels, while maintaining a stable position relative to the SCT. As motile cells that navigate the stromal space in chorionic villi between the foetal and maternal vasculature, HBCs are thought to play a role in mediating the transport of nutrients and maternal antibodies across that space ^40,41,46^. The implication of this spatial repositioning of HBCs in placentas with past malaria needs further work.

Taken together, our results suggest that even though there are residual effects from malaria on critical foetal outcomes such as birthweight, any activated HBCs have either been replaced or returned to a basal state. However, we did find evidence that infections early in pregnancy could involve a disruption of the normal vascular development of the placenta. The data we collected on the Visium platform suggests that placentas exposed to past malaria infections have a different transcriptomic profile from those where evidence of malaria infection is absent. As such, future studies should now turn their attention to other chorionic villi compartments – such as the SCT, cytotrophoblasts, fibroblasts or the vascular endothelium – to understand their role as markers and/or mediators of these continuing pathological processes resulting from placental injury after malaria infection.

## Meetings

Part of this work was presented at the American Society for Tropical Medicine and Hygiene conference in Seattle, 2022

## Author contributions

**RA:** Conceptualization, Methodology, Formal analysis, Investigation, Data curation, Writing – original draft, Writing - Review & Editing, Visualisation **MD:** Formal analysis, Data curation, Writing - Review & Editing, Visualisation **RH:** Validation, Formal analysis, Data curation **YC:** Formal analysis, Data curation, Writing - Review & Editing, Visualisation **KF:** Writing - Review & Editing **LW:** Software, Formal analysis, Data curation **KLR:** Supervision **CA:** Investigation **LL:** Investigation **PH:** Formal analysis, Data curation **DA-Z:** Validation, Resources **EM:** Investigation, Writing - Review & Editing, Project administration **GMh:** Investigation, Writing - Review & Editing, Project administration **SK:** Investigation **LR:** Writing - Review & Editing **CB:** Writing - Review & Editing **GMz:** Investigation, Writing - Review & Editing, Project administration **MNM:** Investigation, Writing - Review & Editing, Project administration **SB:** Validation, Writing - Review & Editing **KP:** Resources, Writing - Review & Editing, Project administration, Funding acquisition **SRP:** Conceptualization, Resources, Writing - Review & Editing, Supervision, Project administration, Funding acquisition.

## Acknowledgements

The REVAMP trial from which samples derive was funded by the Bill & Melinda Gates Foundation (INV-010612). S-RP is funded by the Australian National Health and Medical Research Council (NHMRC) Fellowships (GNT1158696 and GNT2009047). RA would like to thank Dr. Elizabeth H. Aitken for helpful discussions. We thank WEHI Histology for assistance with FFPE slide preparation and optimising the OPAL staining protocol. We thank the WEHI Genomic facility for assistance with the Visium 10X protocol and Illumina sequencing. We thank the local field workers, participants, and their families for their involvement in the study.

## References

1. Health Organization, W. *World malaria report* 2022. https://www.who.int/teams/global-malaria-programme.

2. Rogerson, S. J., Hviid, L., Duffy, P. E., Leke, R. F. & Taylor, D. W. Malaria in pregnancy: pathogenesis and immunity. Lancet Infect Dis 7, 105– 117 (2007).

3. Bulmer, J. N., Rasheed, F. N., Morrison, L., Francis, N. & Greenwood, B. M. Placental malaria. I. Pathological classification. Histopathology 22, 211–8 (1993).

4. Moore, K. A., Simpson, J. A., Scoullar, M. J. L., McGready, R. & Fowkes, F. J. I. Quantification of the association between malaria in pregnancy and stillbirth: a systematic review and meta-analysis. Lancet Glob Health 5, e1101–e1112 (2017).

5. Guyatt, H. L. & Snow, R. W. Impact of malaria during pregnancy on low birth weight in sub-Saharan Africa. Clin Microbiol Rev 17, 760–9, table of contents (2004).

6. Umbers, A. J., Aitken, E. H. & Rogerson, S. J. Malaria in pregnancy: small babies, big problem. Trends Parasitol 27, 168–75 (2011).

7. World Health Organization. *Guideline WHO Guidelines for malaria - 14 March 2023*. http://apps.who.int/bookorders. (2023).

8. Walker, P. G. T., ter Kuile, F. O., Garske, T., Menendez, C. & Ghani, A. C. Estimated risk of placental infection and low birthweight attributable to Plasmodium falciparum malaria in Africa in 2010: a modelling study. Lancet Glob Health 2, e460–e467 (2014).

9. Maltepe, E., Bakardjiev, A. I. & Fisher, S. J. The placenta: transcriptional, epigenetic, and physiological integration during development. J Clin Invest 120, 1016 (2010).

10. Maltepe, E. & Fisher, S. J. Placenta: The Forgotten Organ. Annu Rev Cell Dev Biol 31, 523–552 (2015).

11. Turco, M. Y. & Moffett, A. Development of the human placenta. Development (Cambridge) vol. 146 Preprint at 10.1242/dev.163428 (2019).

12. Fried, M., Muga, R. O., Misore, A. O. & Duffy, P. E. Malaria Elicits Type 1 Cytokines in the Human Placenta: IFN-γ and TNF-α Associated with Pregnancy Outcomes. The Journal of Immunology 160, (1998).

13. Abrams, E. T. et al. Host response to malaria during pregnancy: placental monocyte recruitment is associated with elevated beta chemokine expression. J Immunol 170, 2759–64 (2003).

14. Lucchi, N. W., Peterson, D. S. & Moore, J. M. Immunologic activation of human syncytiotrophoblast by Plasmodium falciparum. Malar J 7, 42 (2008).

15. Lucchi, N. W. et al. Natural hemozoin stimulates syncytiotrophoblast to secrete chemokines and recruit peripheral blood mononuclear cells. Placenta 32, 579–85 (2011).

16. Ataíde, R., Mayor, A. & Rogerson, S. J. Malaria, primigravidae, and antibodies: Knowledge gained and future perspectives. Trends in Parasitology vol. 30 85–94 Preprint at (2014).

17. Aitken, E. H. et al. Developing a multivariate prediction model of antibody features associated with protection of malaria-infected pregnant women from placental malaria. Elife 10, 1–30 (2021).

18. Tutterrow, Y. L. et al. High levels of antibodies to multiple domains and strains of VAR2CSA correlate with the absence of placental malaria in Cameroonian women living in an area of high Plasmodium falciparum transmission. Infect Immun 80, 1479–90 (2012).

19. Dobano, C. et al. High production of pro-inflammatory cytokines by maternal blood mononuclear cells is associated with reduced maternal malaria but increased cord blood infection. Malar J 17, 177 (2018).

20. Conroy, A. L., Liles, W. C., Molyneux, M. E., Rogerson, S. J. & Kain, K. C. Performance characteristics of combinations of host biomarkers to identify women with occult placental malaria: a case-control study from Malawi. PLoS One 6, e28540 (2011).

21. Conroy, A. et al. C5a enhances dysregulated inflammatory and angiogenic responses to malaria in vitro: potential implications for placental malaria. PLoS One 4, e4953 (2009).

22. Conroy, A. L. et al. Complement Activation and the Resulting Placental Vascular Insufficiency Drives Fetal Growth Restriction Associated with Placental Malaria. Cell Host Microbe 13, 215–226 (2013).

23. Id, R. E. E. et al. Early malaria infection, dysregulation of angiogenesis, metabolism and inflammation across pregnancy, and risk of preterm birth in Malawi: A cohort study. 1–21 (2019) doi:10.1371/journal.pmed.1002914.

24. Ataíde, R. et al. Malaria in Pregnancy Interacts with and Alters the Angiogenic Profiles of the Placenta. PLoS Negl Trop Dis 9, e0003824 (2015).

25. Muehlenbachs, A., Mutabingwa, T. K., Edmonds, S., Fried, M. & Duffy, P. E. Hypertension and Maternal–Fetal Conflict during Placental Malaria. PLoS Med 3, e446 (2006).

26. Kabyemela, E. R. et al. Maternal peripheral blood level of IL-10 as a marker for inflammatory placental malaria. Malar J 7, 26 (2008).

27. Dong, S. et al. CXCL9 response to pregnancy malaria is associated with low birth weight deliveries. Infect Immun (2012) doi:10.1128/IAI.00220-12.

28. Souza, R. M. et al. Placental Histopathological Changes Associated with Plasmodium vivax Infection during Pregnancy. PLoS Negl Trop Dis 7, e2071 (2013).

29. Bulmer, J. N., Rasheed, F. N., Morrison, L., Francis, N. & Greenwood, B. M. Placental malaria. II. A semi-quantitative investigation of the pathological features. Histopathology 22, 219–25 (1993).

30. McGready, R. et al. The effects of Plasmodium falciparum and P. vivax infections on placental histopathology in an area of low malaria transmission. Am J Trop Med Hyg 70, 398–407 (2004).

31. Chaikitgosiyakul, S. et al. A morphometric and histological study of placental malaria shows significant changes to villous architecture in both Plasmodium falciparum and Plasmodium vivax infection. Malar J 13, 4 (2014).

32. Hoeffel, G. & Ginhoux, F. Ontogeny of tissue-resident macrophages. Frontiers in Immunology vol. 6 Preprint at 10.3389/fimmu.2015.00486 (2015).

33. Reyes, L., Wolfe, B. & Golos, T. Hofbauer Cells: Placental Macrophages of Fetal Origin. in Macrophages, Results and Problems in Cell Differentiation (ed. Kloc, M.) vol. 62 45–60 (Springer International Publishing, 2017).

34. Mezouar, S., Katsogiannou, M., Ben Amara, A., Bretelle, F. & Mege, J. L. Placental macrophages: Origin, heterogeneity, function and role in pregnancy-associated infections. Placenta 103, 94–103 (2021).

35. Reyes, L. & Golos, T. G. Hofbauer cells: Their role in healthy and complicated pregnancy. Front Immunol 9, 1–8 (2018).

36. Tang, Z., Abrahams, V. M., Mor, G. & Guller, S. Placental Hofbauer cells and complications of pregnancy. Ann N Y Acad Sci 1221, 103–108 (2011).

37. Young, O. M. et al. Toll-like Receptor-Mediated Responses by Placental Hofbauer Cells (HBCs): A Potential Pro-Inflammatory Role for Fetal M2 Macrophages. American Journal of Reproductive Immunology 73, 22–35 (2015).

38. Seval, Y., Korgun, E. T. & Demir, R. Hofbauer Cells in Early Human Placenta: Possible Implications in Vasculogenesis and Angiogenesis. Placenta 28, 841–845 (2007).

39. Loegl, J. et al. Hofbauer cells of M2a, M2b and M2c polarisation may regulate feto-placental angiogenesis. Reproduction 152, 447–455 (2016).

40. Saji, F., Samejima, Y., Kamiura, S. & Koyama, M. Dynamics of immunoglobulins at the feto-maternal interface. Rev Reprod. 4, 81–9. (1999).

41. Bastin, J., Drakesmith, H., Rees, M., Sargent, I. & Townsend, A. Localisation of proteins of iron metabolism in the human placenta and liver. Br J Haematol 134, 532–543 (2006).

42. Vento-Tormo, R. et al. Single-cell reconstruction of the early maternal–fetal interface in humans. Nature 563, 347–353 (2018).

43. Li, H., Huang, Q., Liu, Y. & Garmire, L. X. Single cell transcriptome research in human placenta. Reproduction 160, R155–R167 (2020).

44. Thomas, J. R. et al. Phenotypic and functional characterisation of first-trimester human placental macrophages, Hofbauer cells. Journal of Experimental Medicine 218, (2020).

45. Caesarine, A., et al. Single Cell Profiling of Hofbauer Cells and Fetal Brain Microglia Reveals Shared Programs and Functions. BioRxiv (2021) doi:10.2139/ssrn.3985607.

46. Zulu, M. Z., Martinez, F. O., Gordon, S. & Gray, C. M. The Elusive Role of Placental Macrophages: The Hofbauer Cell. Journal of Innate Immunity Preprint at 10.1159/000497416 (2019).

47. Fakonti, G., Pantazi, P., Bokun, V. & Holder, B. Placental Macrophage (Hofbauer Cell) Responses to Infection During Pregnancy : A Systematic Scoping Review. 12, 1–20 (2022).

48. Sisino, G. et al. Diabetes during pregnancy influences Hofbauer cells, a subtype of placental macrophages, to acquire a pro-inflammatory phenotype. Biochim Biophys Acta Mol Basis Dis 1832, 1959–1968 (2013).

49. Azari, S. et al. Hofbauer Cells Spread Listeria monocytogenes among Placental Cells and Undergo Pro-Inflammatory Reprogramming while Retaining Production of Tolerogenic Factors. https://doi.org/10 (2021).

50. Schliefsteiner, C. et al. Human placental Hofbauer cells maintain an anti-inflammatory M2 phenotype despite the presence of gestational diabetes mellitus. Front Immunol 8, 1–17 (2017).

51. Schwartz, D. A. Viral infection, proliferation, and hyperplasia of Hofbauer cells and absence of inflammation characterise the placental pathology of fetuses with congenital Zika virus infection. Arch Gynecol Obstet 1–8 (2017) doi:10.1007/s00404-017-4361-5.

52. Morotti, D. et al. Molecular pathology analysis of sars-cov-2 in syncytiotrophoblast and hofbauer cells in placenta from a pregnant woman and fetus with covid-19. Pathogens 10, (2021).

53. Schwartz, D. A. et al. Hofbauer cells and COVID-19 in pregnancy: Molecular pathology analysis of villous macrophages, endothelial cells, and placental findings from 22 placentas infected by SARS-CoV-2 with and without fetal transmission. Arch Pathol Lab Med 145, 1328–1340 (2021).

54. Johnson, E. L. & Chakraborty, R. Placental Hofbauer cells limit HIV-1 replication and potentially offset mother to child transmission (MTCT) by induction of immunoregulatory cytokines. Retrovirology 9, 101 (2012).

55. Johnson, E. L., Chu, H., Byrareddy, S. N., Spearman, P. & Chakraborty, R. Placental Hofbauer cells assemble and sequester HIV-1 in tetraspanin-positive compartments that are accessible to broadly neutralising antibodies. J Int AIDS Soc 18, 19385 (2015).

56. Kim, J.-S. et al. Involvement of Hofbauer cells and maternal T cells in villitis of unknown aetiology. Histopathology 52, 457–64 (2008).

57. Grigoriadis, C. et al. Hofbauer cells morphology and density in placentas from normal and pathological gestations. Revista Brasileira de Ginecologia e Obstetrícia 35, 407–412 (2013).

58. Gaw, S. L. et al. Differential Activation of Fetal Hofbauer Cells in Primigravidas Is Associated with Decreased Birth Weight in Symptomatic Placental Malaria. Malar Res Treat 2019, 1–10 (2019).

59. Tkachuk, A. N. et al. Malaria enhances expression of CC chemokine receptor 5 on placental macrophages. J Infect Dis 183, 967–72 (2001).

60. Pasricha, S.-R. et al. Ferric carboxymaltose versus standard-of-care oral iron to treat second-trimester anaemia in Malawian pregnant women: a randomised controlled trial. Lancet 401, 1595–1609 (2023).

61. Mwangi, M. N. et al. Protocol for a multicentre, parallel-group, open-label randomised controlled trial comparing ferric carboxymaltose with the standard of care in anaemic Malawian pregnant women: The REVAMP trial. BMJ Open 11, (2021).

62. Schindelin, J., et al. Fiji: An open-source platform for biological-image analysis. Nature Methods vol. 9 676–682 Preprint at 10.1038/nmeth.2019 (2012).

63. Berg, S. et al. ilastik: interactive machine learning for (bio)image analysis. Nat Methods 16, 1226–1232 (2019).

64. Weigert, M., Schmidt, U., Haase, R., Sugawara, K. & Myers, G. Star-convex Polyhedra for 3D Object Detection and Segmentation in Microscopy. in 2020 IEEE Winter Conference on Applications of Computer Vision (WACV) 3655–3662 (IEEE, 2020). doi:10.1109/WACV45572.2020.9093435.

65. Schmidt, U., Weigert, M., Broaddus, C. & Myers, G. Cell Detection with Star-Convex Polygons. in Medical Image Computing and Computer Assisted Intervention - {MICCAI} 21st International Conference 265–273 (2018). doi:10.1007/978-3-030-00934-2_30.

66. Swieboda, D. et al. Baby’s First Macrophage: Temporal Regulation of Hofbauer Cell Phenotype Influences Ligand-Mediated Innate Immune Responses across Gestation. The Journal of Immunology 204, 2380–2391 (2020).

67. Tang, P. M.-K. et al. Neural transcription factor Pou4f1 promotes renal fibrosis via macrophage–myofibroblast transition. Proceedings of the National Academy of Sciences 117, 20741–20752 (2020).

68. Meng, X. M. et al. Inflammatory macrophages can transdifferentiate into myofibroblasts during renal fibrosis. Cell Death Dis 7, (2016).

69. Abu El-Asrar, A. M., et al. Macrophage-Myofibroblast Transition Contributes to Myofibroblast Formation in Proliferative Vitreoretinal Disorders. Int J Mol Sci 24, (2023).

70. Kim, S. S. et al. Coexpression of myofibroblast and macrophage markers: Novel evidence for an in vivo plasticity of chorioamniotic mesodermal cells of the human placenta. Laboratory Investigation 88, 365– 374 (2008).

71. Chambers, M., et al. Macrophage Plasticity in Reproduction and Environmental Influences on Their Function. Frontiers in Immunology vol. 11 Preprint at 10.3389/fimmu.2020.607328 (2021).

72. Ajami, B., Bennett, J. L., Krieger, C., Tetzlaff, W. & Rossi, F. M. V. Local self-renewal can sustain CNS microglia maintenance and function throughout adult life. Nat Neurosci 10, 1538–1543 (2007).

73. Rosenberg, A. Z., Weiying, Y., Hill, D. A., Reyes, C. A. & Schwartz, D. A. Placental pathology of zika virus: Viral infection of the placenta induces villous stromal macrophage (Hofbauer Cell) proliferation and hyperplasia. Arch Pathol Lab Med 141, 43–48 (2017).

74. Baergen, R. N. Manual of Pathology of the Human Placenta. (Springer US, 2011). doi:10.1007/978-1-4419-7494-5.

75. Pearson, R. D. Placental Malaria: Hypertension, VEGF, and Prolactin. PLoS Med 4, 2 (2007).

76. Silver, K. L., Zhong, K., Leke, R. G. F., Taylor, D. W. & Kain, K. C. Dysregulation of Angiopoietins Is Associated with Placental Malaria and Low Birth Weight. PLoS One 5, e9481 (2010).

