## Supplementary information for "Spatial analysis of Hofbauer cell transcriptome, distribution and morphology in placentas exposed to *Plasmodium falciparum*"

**Full title:** Spatial analysis of Hofbauer cell transcriptome, distribution and morphology in placentas exposed to *Plasmodium falciparum*. **– SUPPLEMENTARY INFORMATION.**

Ricardo Ataide[1,2,*], Rebecca Harding[1], Malindrie Dharmaratne[1,3,4], Yunshun Chen[1,3,4], Katherine Fielding[1], Lachlan Whitehead[3,5], Kelly L. Rogers[3,5], Casey Anttilla[3,6], Ling Ling[6], Peter Hickey[3,6], Daniela Amann-Zalcenstein[3,6], Ernest Moya[7,8], Gomezghani Mhango[7], Steve Kamiza[9], Louise Randall[1], Cavan Bennett[1,3], Glory Mzembe[7,8], Martin N. Mwangi[7,8,10,11], Sabine Braat[1,12], Kamija Phiri[7,8], Sant-Rayn Pasricha[1,13,14].

[1] Population Health and Immunity division, The Walter and Eliza Hall Institute, Parkville, VIC, Australia

[2] Department of Infectious Diseases, The Peter Doherty Institute for Infection and Immunity, Melbourne, VIC, Australia

[3] Department of Medical Biology, University of Melbourne, Melbourne, VIC, Australia

[4] ACRF Cancer Biology and Stem Cells Division, The Walter and Eliza Hall Institute of Medical Research, Parkville, VIC 3052, Australia

[5] Centre for Dynamic Imaging, The Walter and Eliza Hall Institute, Parkville, VIC, Australia.

[6] Advanced Genomics Facility, The Walter and Eliza Hall Institute, Parkville, VIC, Australia.

[7] Training and Research Unit of Excellence (TRUE), Chichiri, Blantyre, Malawi

[8] School of Global and Public Health, Kamuzu University of Health Sciences, Chichiri, Blantyre, Malawi

[9] Department of Pathology, Kamuzu University of Health Sciences, Chichiri, Blantyre, Malawi.

[10] Healthy Mothers Healthy Babies Consortium, Micronutrient Forum, Washington DC, USA

[11] Division of Human Nutrition, Wageningen University, Wageningen, The Netherlands

[12] Centre for Epidemiology and Biostatistics, Melbourne School of Population and Global Health, The University of Melbourne, Melbourne, VIC, Australia

[13] Diagnostic Hematology, The Royal Melbourne Hospital, Parkville, Australia

[14] Clinical Hematology at the Royal Melbourne Hospital and Peter MacCallum Cancer Centre, Parkville, Australia

Figure legends

**Supplementary figure 1- Visium 10X: assay reliability and differentially expressed genes across all placental samples**.

Two independent Visium Spatial for FFPE Gene Expression Kit, Human Transcriptome slides containing 4 capture areas each (8 tissues in total) were pooled and sequenced on the Illumina NextSeq2000 according to 10X Genomics' recommendations and heatmaps of differentially expressed genes were constructed. (A) A commercially acquired placental sample from Cureline, Inc., CA, USA and a Malawian sample from our trials were used on both slides to control for tissue quality and assay reliability (identified as Dup, under the heatmap). In total, seven tissues had enough quality to generate sequencing data (one tissue lifted off the slide and did not allow for any sequencing to be performed). Both Dup samples had highly similar sequencing results across the two independent runs. All Malawian samples were more similar to each other than to the Cureline placental tissue sample.

**Supplementary figure 2- GeoMX assay quality control and sensitivity**

(A) Graph representing the sequencing saturation achieved. No samples were below the 50% warning where the counts become less reliable. Sensitivity for low expressing genes is depressed with lower sequencing saturation. (B) Q3 normalization was used as it consistently outperforms other methods like Negative Probe and Housekeeper Normalization. (C) Number of genes expressed above the limit of quantitation - Geometric Mean of the Negative probes, multiplied by the Geometric Standard Deviation to the second power.

**Supplementary figure 3- KEGG pathway analysis using Visium data**

KEGG Pathway analysis was used to predict the networks with differentially expressed genes between Malawian placental samples with or without evidence of malaria infection. Analysis was done in R using the 'kegga' function and setting the FDR threshold to 0.2.

**Supplementary figure 4- Placental cell types identified in GeoMX data using 'Visium deconvolution' gene sets**

The GeoMX Q3-normalization counts for gene sets used to deconvolute the Visium data are plotted. Each gene set is shown above the cell type it identifies. Placental samples have been grouped by malaria infection status (Absent, Past and Active).

**Supplementary figure 5 – Hypothesis driven gene lists**

#### Supplementary figure 1

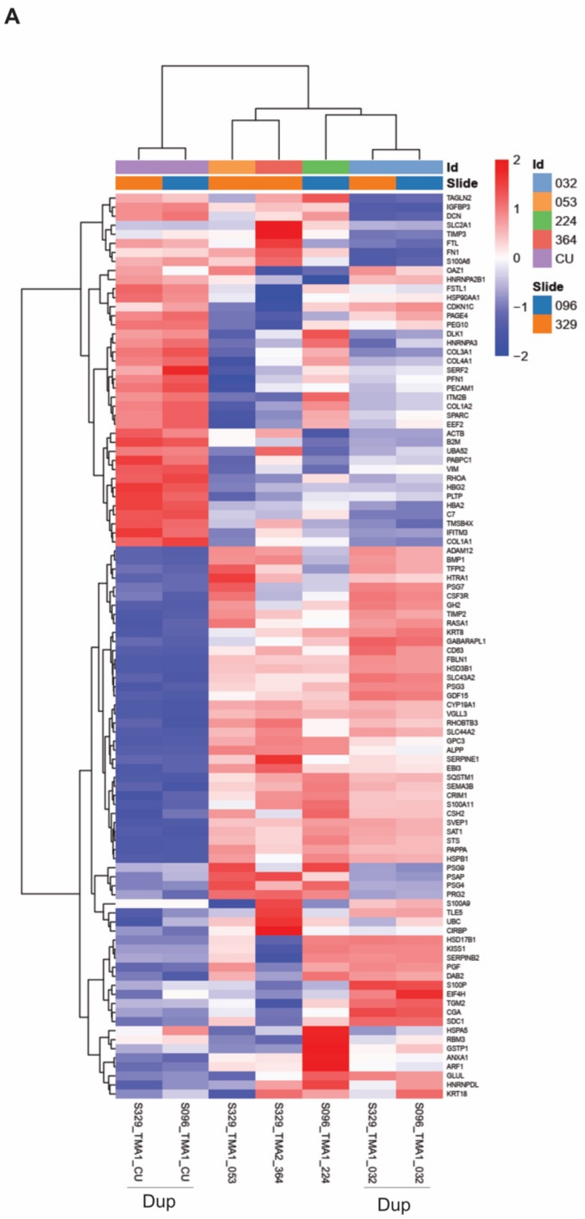

### Supplementary figure 2

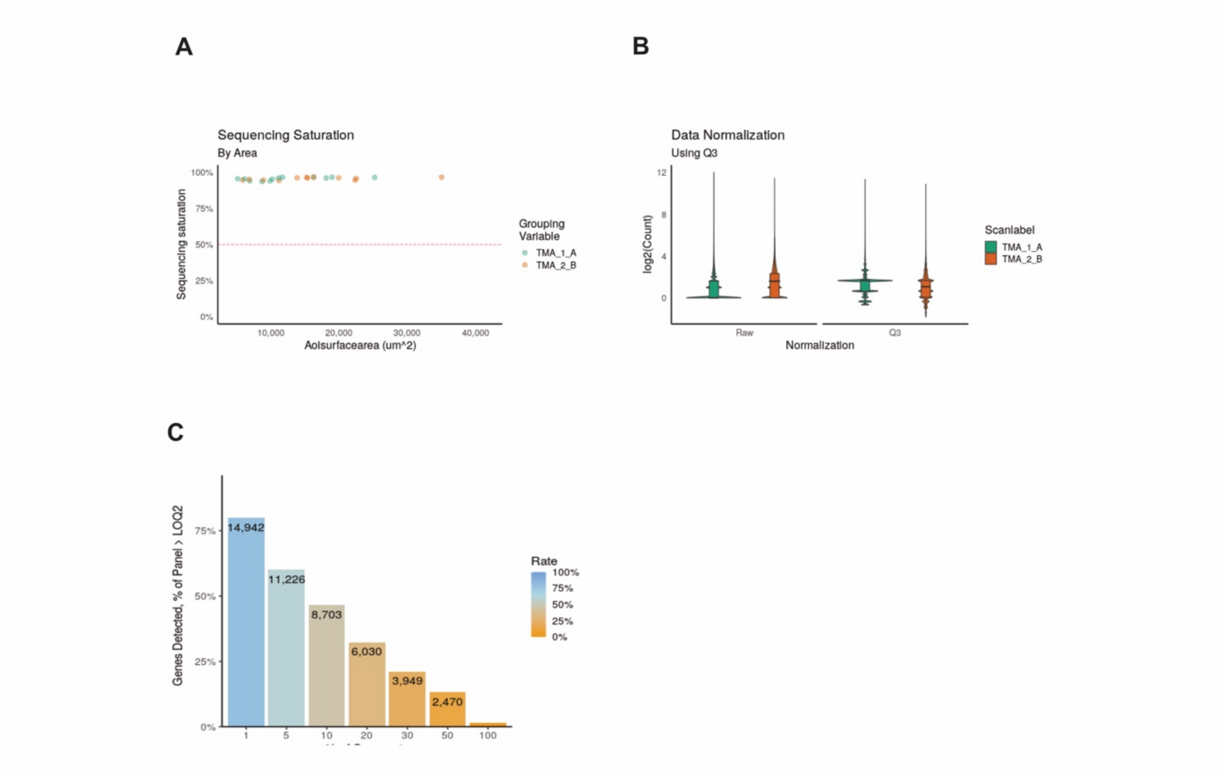

#### Supplementary figure 3

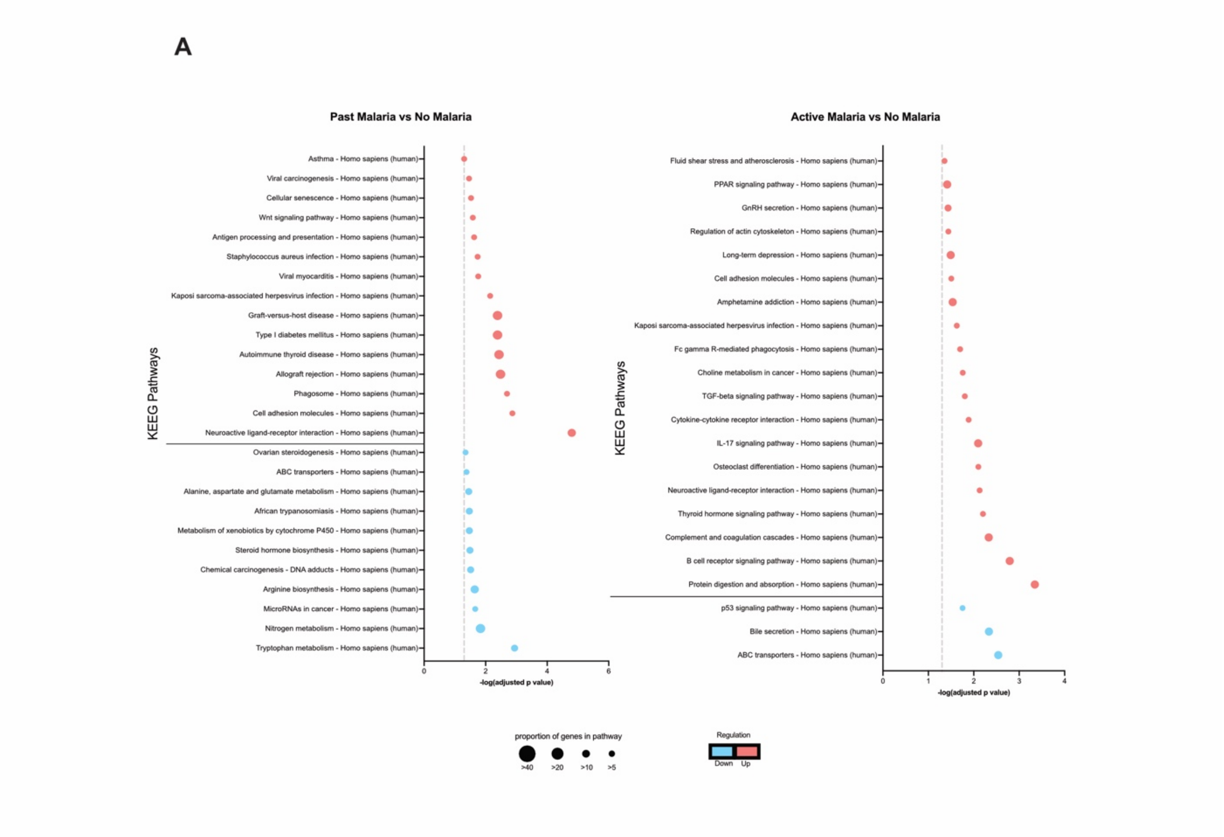

#### Supplementary figure 4

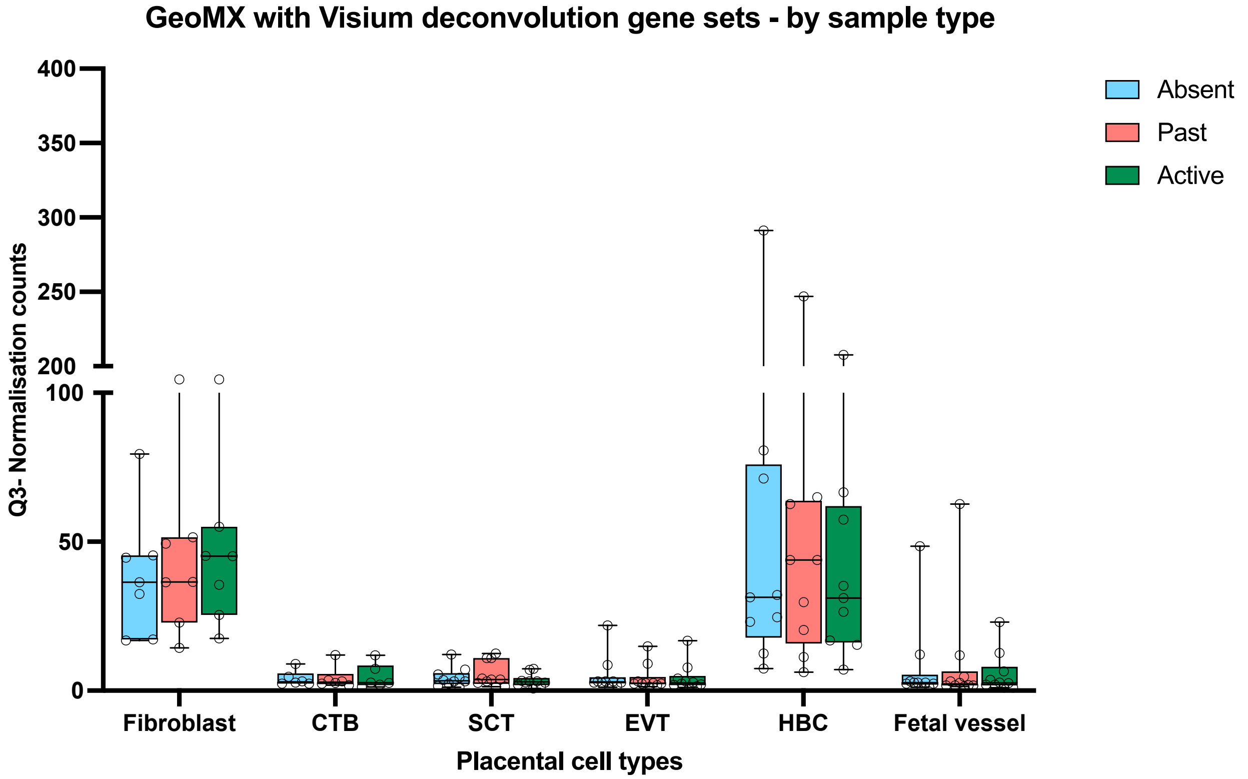

#### Supplementary figure 5

**Adhesion and Angiogenesis**

| | |
|---|---|
| SLC2A1 | VEGFA |
| SLC2A4 | VEGFB |
| NOS1 | ANGPTL1 |
| NOS2 | FLT1 |
| NOS3 | FLT3 |
| MKI67 | FLT4 |
| ICAM1 | ANGPT1 |
| ICAM2 | ANGPT2 |
| ICAM3 | MTOR |
| ICAM4 | TIE1 |
| ICAM5 | HIF1A |
| VCAM1 | |

**Iron response and regulation**

| | | |
|---|---|---|
| FTH1 | SLC40A1 | BMP1 |
| FTMT | HJV | BMP2 |
| HFE | TFRC | BMP3 |
| IREB2 | SLC11A2 | BMP5 |
| HAMP | SLC39A8 | BMP6 |
| ACO1 | SLC39A14 | BMP7 |
| CP | STEAP4 | SMAD1 |
| TF | HAVCR1 | SMAD4 |
| TMPRSS6 | CXCR4 | SMAD5 |
| SCARA5 | HMOX1 | SMAD6 |
| | JAK1 | SMAD7 |
| | JAK2 | SMAD2 |
| | STAT1 | |
| | STAT5A | |

**Macrophage activation**

| | | |
|---|---|---|
| FCGR1A | CXCL1 | TLR1 |
| FCGR2B | CXCL2 | TLR2 |
| FCGR2C | CXCL5 | TLR3 |
| FCGR3A | CXCL6 | TLR4 |
| FCGRT | CXCL9 | TLR5 |
| CD36 | CXCL8 | TLR6 |
| CD80 | CXCL10 | TLR7 |
| CD86 | CXCL11 | TLR8 |
| CD40 | CXCL12 | TLR9 |
| IL1B | CXCL13 | TLR10 |
| IL12B | CXCL14 | |
| IL6 | CXCL16 | |
| IL10 | CXCL17 | |
| IL13 | | |
| IL18 | | |
| IFNG | | |
| TGFB1 | | |

Supplementary Table 1 – Clinical association with birth weight

| | | Unadjusted regression | | | Adjusted regression* | | |
|---|---|---|---|---|---|---|---|
| | Birth weight n, Mean (SD) | Mean difference | 95% Confidence Interval | p-value | Mean difference | 95% Confidence Interval | p-value |
| Placental malaria | | | | | | | |
| Absent[a] | 391, 2937.2 (522.2) | Reference | | | | | |
| Present (Past[b]/Active[c]) | 319, Mean (SD) | -119.3 | -202.5, -36.1 | 0.005 | | | |
| Placental malaria | | | | | | | |
| Absent[a] | 391, 2937.2 (522.2) | Reference | | | Reference | | |
| Past[b] | 192, 2814.8 (467.3) | -123.0 | -209.9, -36.1 | 0.006 | -80.9 | -165.9, -3.7 | 0.040 |
| Active[c] | 27, 2844.8 (432.5) | -93.0 | - 288.9, 103.0 | 0.352 | -9.7 | -174.3, 154.8 | 0.943 |

SD: Standard Deviation.

[*] Model adjusted for gravidity (primigravidae, secundigravidae, multigravidae), maternal HIV status (negative, positive), maternal anaemia at delivery (yes, no), maternal iron-deficiency at delivery (yes, no), gestational age at delivery (continuous [weeks]), infant sex (female, male) and trial arm (IV iron or Oral iron) to obtain the direct effect of placental malaria on birthweight.

[a] Absent is defined as absence of tissue evidence for malaria (parasites or pigment). N=339 in the adjusted model.

[b] Past malaria includes only those women where malaria pigment was detected in the placenta tissue in the absence of any circulating parasites. N=173 in the adjusted model.

[c] Active malaria includes tissues where we detected the presence of circulating parasites in the intervillous space. Twenty placental tissues had evidence of pigment in the tissue in addition to circulating parasites, and thus could have been further classified as Chronic infections. N=27 in the adjusted model.

Supplementary Table 2 – Association with immunofluorescence-derived measurements

| | | Unadjusted regression | | | Adjusted regression* | | |
|---|---|---|---|---|---|---|---|
| | Fetal vessel proportion n, Mean (SD) | Mean difference | 95% Confidence Interval | p-value | Mean difference | 95% Confidence Interval | p-value |
| Placental malaria | | | | | | | |
| Absent[a] | 54, 59.2 (45.4) | Reference | | | Reference | | |
| Past[b] | 47, 36.4 (29.5) | -22.8 | -38.1, -7.6 | 0.004 | -18.3 | -36.6, -0.1 | 0.049 |
| Active[c] | 4, 37.0 (29.1) | -22.2 | -61.9, 17.4 | 0.268 | -25.7 | -65.0, 13.6 | 0.197 |
| Placental malaria | Distance of HBC to foetal vessel (pixels) n, Mean, (SD) | Mean difference | 95% Confidence Interval | p-value | Mean difference | 95% Confidence Interval | p-value |
| Absent[a] | 55, 6.7 (6.9) | Reference | | | Reference | | |
| Past[b] | 47, 23.2 (35.5) | 16.5 | 6.9, 26.0 | 0.001 | 15.8 | 3.3, 28.4 | 0.014 |
| Active[c] | 4, 3.3 (1.9) | -3.5 | -28.4, 21.5 | 0.784 | -6.7 | -34.5, 21.2 | 0.635 |
| | Proportion of Ki67+ HBC n, Mean (SD) | Mean difference | 95% Confidence Interval | p-value | Mean difference | 95% Confidence Interval | p-value |
| Placental malaria | | | | | | | |
| Absent[a] | 56, 0.7 (0.9) | Reference | | | Reference | | |
| Past[b] | 48, 0.5 (0.7) | -0.1 | -0.5, 0.2 | 0.389 | -0.2 | -0.6, 0.3 | 0.414 |
| Active[c] | 5, 2.0 (2.0) | 1.3 | 0.5, 2.2 | 0.001 | 1.4 | 0.5, 2.3 | 0.002 |

SD: Standard Deviation.

* Model adjusted for gravidity (primigravidae, secundigravidae, multigravidae), maternal HIV status (negative, positive), maternal anaemia at delivery (yes, no), maternal iron-deficiency at delivery (yes, no), gestational age at delivery (continuous [weeks]), infant sex (female, male) and trial arm (IV iron or Oral iron) to obtain the direct effect of placental malaria on birthweight.

[a] Absent is defined as absence of tissue evidence for malaria (parasites or pigment).

[b] Past malaria includes only those women where malaria pigment was detected in the placenta tissue in the absence of any circulating parasites.

[c] Active malaria includes tissues where we detected the presence of circulating parasites in the intervillous space. Twenty placental tissues had evidence of pigment in the tissue in addition to circulating parasites, and thus could have been further classified as Chronic infections.

#### Supplementary methods

*Spatial transcriptomics – Detailed Visium slide preparation and sequencing*

For 10X Genomics, Visium Spatial for FFPE Gene Expression Kit, Human Transcriptome, 4 reactions (#1000337, 10x Genomics) were used according to the manufacturer's recommendations. Briefly, 5-μm sections were cut from the TMA and transferred onto a petri dish wrapped in foil. The tissues of interest were dissected from the sections and transferred onto a 42°C water bath. Individual tissue sections were placed onto a Visium Spatial Gene Expression Slide. Each area contained ~5,000 detection spots of 55 μm in diameter and a 100 μm centre-to-centre distance. Each slide included a placental sample from a placenta originating from the United States (purchased from Cureline, Inc., CA, USA) to serve as a control for the quality of the tissue fixation and processing conducted in Malawi and to provide an inter-reliability control for the Visium reactions. Slides were heated at 42°C for 3 hrs on a thermocycler with a Visium PCR Adaptor. Slides were placed in the desiccator at room temperature from overnight up to one week. The Visium slides were processed according to the Visium Spatial Gene Expression for FFPE – Deparaffinization, H&E Staining, Imaging & Decrosslinking protocol, and followed by the Visium Spatial Gene Expression Reagent Kits for FFPE protocol in accordance with the manufacturer's instructions. Gene expression libraries were pooled and sequenced on the Illumina NextSeq2000 (machine details) according to the 10X recommendations. In placental TMAs where the consecutive FFPE sections exposed areas outside of the chorionic villi or where there was carryover from one placental sample to another, the analysis of the results was restricted to a manually selected area of chorionic villi belonging to one placental sample only (see Figure 2A for an example).

Illumina output from 10x Visium sequencing was processed using spaceranger 2.0.0. The demultiplexed spatial transcriptomic data from spaceranger were input to R for preprocessing and downstream analysis. Downstream analysis of the Visium data were restricted to spots within the placental villi that were enriched for HBCs, defined as spots with non-zero *LYVE1* counts. Read counts of *LYVE1* positive spots were aggregated for each Visium slide, creating pseudobulk samples similar to RNA-seq data. Lowly expressed genes were filtered using the filterByExpr function in edgeR (v3.41.3) followed by TMM normalization. The edgeR quasi-likelihood pipeline was used to identify differentially expressed genes between No Malaria, Past Malaria and Active Malaria groups.

*Chromogenic immunohistochemistry and Opal Polaris multiplexed immunofluorescence*

All antibodies used for multiplexed immunofluorescence were validated individually first by chromogenic singleplex immunohistochemistry (IHC) and then by singleplex immunofluorescence using the Opal 7 kit reagents (NEL871001KT, Akoya Biosciences, Waltham, MA, USA). In summary, for IHC, 5-μm sections of FFPE tissue were dewaxed and rehydrated following Xylene immersion and decreasing Ethanol concentrations. All slides were left in dd$H_2O$ overnight. Antigen retrieval was achieved with Antigen Retrieval solution, pH 9 from DAKO (S236784-2), for 20 min. Endogenous peroxidase activity blocking and non-specific protein blocking was achieved after 10 min with DAKO dual endogenous peroxidase block (S2003) and Blocking buffer (5% goat serum + 1% BSA in Tris-Buffered Saline – Tween 0.05% (TBST)), respectively. All antibodies were diluted in a 1:1 TBST:Blocking buffer prior to use. Incubations were 30 min long, followed by 1:300 dilution of Biotinylated Goat Anti-Mouse IgG Antibody (BA-9200, VECTORLAB®).

Visualisation was achieved with VECTASTAIN® Elite ABC-HRP Reagent, Peroxidase (PK-7100) for 30 min followed by ImmPACT® DAB Substrate, Peroxidase (HRP) (SK-4105). Slides were counterstained with Haemotoxylin and coverslipped before being digitally scanned. For Opal multiplex immunostaining, 2mm core TMAs were prepared after manual selection of areas of Chorionic villi. Opal 7 kit (NEL871001KT, Akoya Biosciences, Waltham, MA, USA) were used for the staining following standard Opal protocols. All slides were scanned on a PhenoImager-HT with a 0.45 NA objective. Primary antibodies and concentrations used:

| Description | Company | Product number | Optimised dilution |
|---|---|---|---|
| CD163 (Ms x Hu) - clone 10D6 | Thermofisher | MA5-11458 | 1:50 |
| Cytokeratin-7 (Ms x Hu) - Clone OV-TL 12/30 | Invitrogen | MA5-11986 | 1:1000 |
| CD31 (Ms x Hu) - Clone JC/70A | ABCAM | ab9498 | 1:200 |
| HIF1-alpha (Rb x Hu) - Clone RPR16897 | ABCAM | ab51608 | 1:200 |
| HLADR (Rb x Hu) - clone EPR3692 | ABCAM | ab92511 | 1:50 |
| Ki67 (Rb x Hu) - clone [SP6] | ABCAM | ab16667 | 1:125 |
